# Multiple methods of diet assessment reveal differences in Atlantic puffin adult and chick diets both between and within years

**DOI:** 10.1101/2024.04.01.587614

**Authors:** William L. Kennerley, Gemma V. Clucas, Donald E. Lyons

## Abstract

Atlantic puffin (*Fratercula arctica*, hereafter “puffin”) reproductive success in the Gulf of Maine (GoM) has decreased following a recent oceanographic regime shift and subsequent rapid warming. Concurrent changes in both the regional forage fish community and puffin chick diet and provisioning rates suggest that inadequate prey resources may be driving this decline. To determine what prey GoM puffins were feeding on during two years of marine heatwave conditions, we assessed puffin diet using two methods: traditional, observational methods that utilize bill-load photography and emerging methods employing fecal DNA metabarcoding. We identified a strong correlation between the composition of chick diet as estimated through traditional and emerging methods, supporting the interpretation of DNA relative read abundance as a quantitative metric of diet composition. Both methods identified the same dominant prey groups, but metabarcoding identified a greater number of species and offered higher taxonomic resolution. Puffin adults and chicks fed on many of the same prey types, although adults consumed a greater variety of taxa and consumed more low quality prey than they provisioned chicks, as predicted by optimal foraging theory. For both age classes, diet varied both between and within years, likely reflecting changes in the local forage fish community in response to environmental variability. During these two years of marine heatwave conditions, puffins exploited unusual abundances of typically-uncommon prey, yet low puffin productivity suggests the observed dietary plasticity was not able to compensate for apparent prey shortages. Continued refinement of molecular tools and the interpretation of the data they provide will enable better assessments of how seabirds of diverse ages and breeding stages are compensating for changing forage fish communities in response to global climate change.

## 1 Introduction

Seabird breeding success is closely related to the availability of adequate prey resources (Cury et al., 2011). As central place foragers, the geographic extent of a breeding seabird’s foraging activity is restricted by the need to return regularly to the nest site (Orians and Pearson, 1979). Thus, while seabirds may spend most of the year capable of traveling vast distances to feed where prey resources and foraging conditions are favorable (Gulka et al., 2023), during the breeding period they become dependent on the prey available within foraging range of the colony. This restriction makes breeding seabirds vulnerable to changes in the distribution, abundance, and composition of local prey resources (Boyd et al., 2017; Fayet et al., 2021).

In response to anthropogenic warming, many marine species have shifted their spatiotemporal patterns of occurrence, including numerous species that are important prey for breeding seabirds (Pinsky et al., 2013; Kleisner et al., 2017; Suca et al., 2021). These prey species, being largely heterothermic, are affected by ocean warming through direct physiological impacts; in contrast, seabirds react to ocean warming indirectly, reacting in response to the changing availability and distribution of their prey (Sydeman et al., 2015). Shifts in marine species distributions may be accelerated by marine heatwaves (MHWs) that temporarily - but often rapidly - bring anomalously warm conditions to a region, altering the distribution of thermal habitat (Lonhart et al., 2019; Jacox et al., 2020). Although changes to a region’s marine community in response to warming are typically two-fold, involving both the “loss” of cold-water adapted species and the “gain” of warm-water adapted species (McLean et al., 2021), not all marine species are equally-suitable as prey for breeding seabirds (Harris et al., 2007; Smith and Craig, 2023). When changing ocean conditions lead to the rapid increase of less-suitable prey taxa, there can be negative impacts on local seabird populations (Descamps et al., 2022).

The suitability of marine taxa as prey for a particular seabird depends on varied traits (e.g., morphology, size, caloric value, lipid-content) that collectively summarize prey quality. When high-quality prey items predominate, seabirds can deliver more energy to chicks per foraging trip and breeding success tends to be high (Schrimpf et al., 2012). In contrast, the increased consumption of low-quality prey types may limit the reproductive success of marine predators (the junk food hypothesis; Österblom et al., 2008; Romano et al., 2006). Although seabirds may be able to compensate for variable prey conditions through behavioral adjustments like increasing foraging effort (Schrimpf et al., 2012), significant declines in the quality of available prey may exceed seabirds’ abilities to compensate (Wanless et al., 2005, 2023; Scopel et al., 2019).

The changing composition of Atlantic puffin (*Fratercula arctica*, hereafter puffins) chick diet has been proposed as the cause of recent declines in puffin reproductive success within the Gulf of Maine (GoM; Kress et al., 2016). A recent (2009-2010) thermal regime shift in the GoM was influenced by above-average ocean temperatures and decreases in zooplankton biomass, with cascading impacts across trophic levels (Johnson et al., 2018; Friedland et al., 2020b; Seidov et al., 2021). The timing of this regime shift closely aligns with both the observed decline in puffin reproductive success and changes in the composition of puffin chick diet (Kress et al., 2016; Scopel et al., 2019; Major et al., 2021). Concerningly, ocean temperatures within the GoM have continued to rise and the region is increasingly affected by MHWs (Pershing et al., 2018; Seidov et al., 2021). The result is that the GoM is now one of the most rapidly-warming parts of the global ocean, with significant declines predicted for many of the region’s species (Pershing et al., 2015, 2021).

When diet monitoring of GoM puffins began in the 1990s, Atlantic herring (*Clupea harengus*, hereafter herring) was among the most commonly observed prey species delivered (Diamond and Devlin, 2003; Diamond, 2021). Herring is an energy-dense species with a high lipid content, making it an important food item for many marine predators within the GoM (Lawson et al., 1998; Overholtz and Link, 2007; Spitz et al., 2010). However, both regional herring recruitment and occurrence in puffin diet have declined markedly in recent decades (Diamond and Devlin, 2003; Kress et al., 2016; Northeast Fisheries Science Center, 2018). In response, puffins have attempted to replace herring by provisioning chicks with alternative prey taxa like haddock (*Meleanogrammus aeglefinnus*) and rough scad (*Trachurus lathami*), neither of which were observed in GoM puffin diet before 2009. Despite the recent inclusion of these species into puffin diet, declining reproductive metrics suggest that these alternative prey species may not be adequate replacements (Scopel et al., 2019).

Puffin diet in the GoM has largely been assessed through the use of bill-load photography to identify the prey items provisioned by adults to chicks at the breeding colony (Kress et al., 2016; Scopel et al., 2019). This method is noninvasive and highly-effective at identifying major prey groups, although visual identification of prey is often limited to low-resolution taxonomic assignments (Gaglio et al., 2017). Furthermore, while bill-load photography may be suitable for estimating chick diet, it is unable to determine the diet of adults since adult puffins consume their prey at sea (Harris and Wanless, 2012). Evidence that seabird chick diet can be used as an effective proxy for adult diet is mixed (Wilson et al., 2004; Bowser et al., 2013) and the diets of adult seabirds, generally, are poorly understood compared to those of chicks (Barrett et al., 2007). However, since adult body condition is known to directly impact reproductive success, adult diet during the breeding period is likely equally as important to understand as chick diet (Erikstad et al., 1997; Barrett et al., 2012). Given the recent decline in GoM puffin reproductive success and continued climate-mediated shifts in the local forage fish community, reassessments of puffin diet and the methods used to determine this are warranted.

In this study, we aim to determine the diet of breeding puffins through the use of two complementary methods: traditional bill-load photography and molecular diet assessment using fecal DNA metabarcoding. From these two methods, we describe the prey types occurring in the diet of puffin chicks and breeding adults across incubation and chick-rearing during two anomalously warm breeding periods. We then examine differences in the composition of diets as determined through the different methodologies as well as between different age classes, breeding stages, and years.

## 2 Methods

### 2.1 Data Collection

#### 2.1.1 Location and Timing

Research occurred at the puffin colony on Matinicus Rock (43.78° N, 68.85° W), an 8-hectare island located 37 km offshore of Rockland, Maine (Figure 1). Matinicus Rock is a part of Maine Coastal Islands National Wildlife Refuge and is cooperatively-managed by the U.S. Fish and Wildlife Service and National Audubon Society’s Seabird Institute. Data collection occurred from June 9^th^ to August 11^th^, 2021 and from May 25^th^ to August 8^th^, 2022, spanning the incubation and chick-rearing periods.

**Figure 1.**
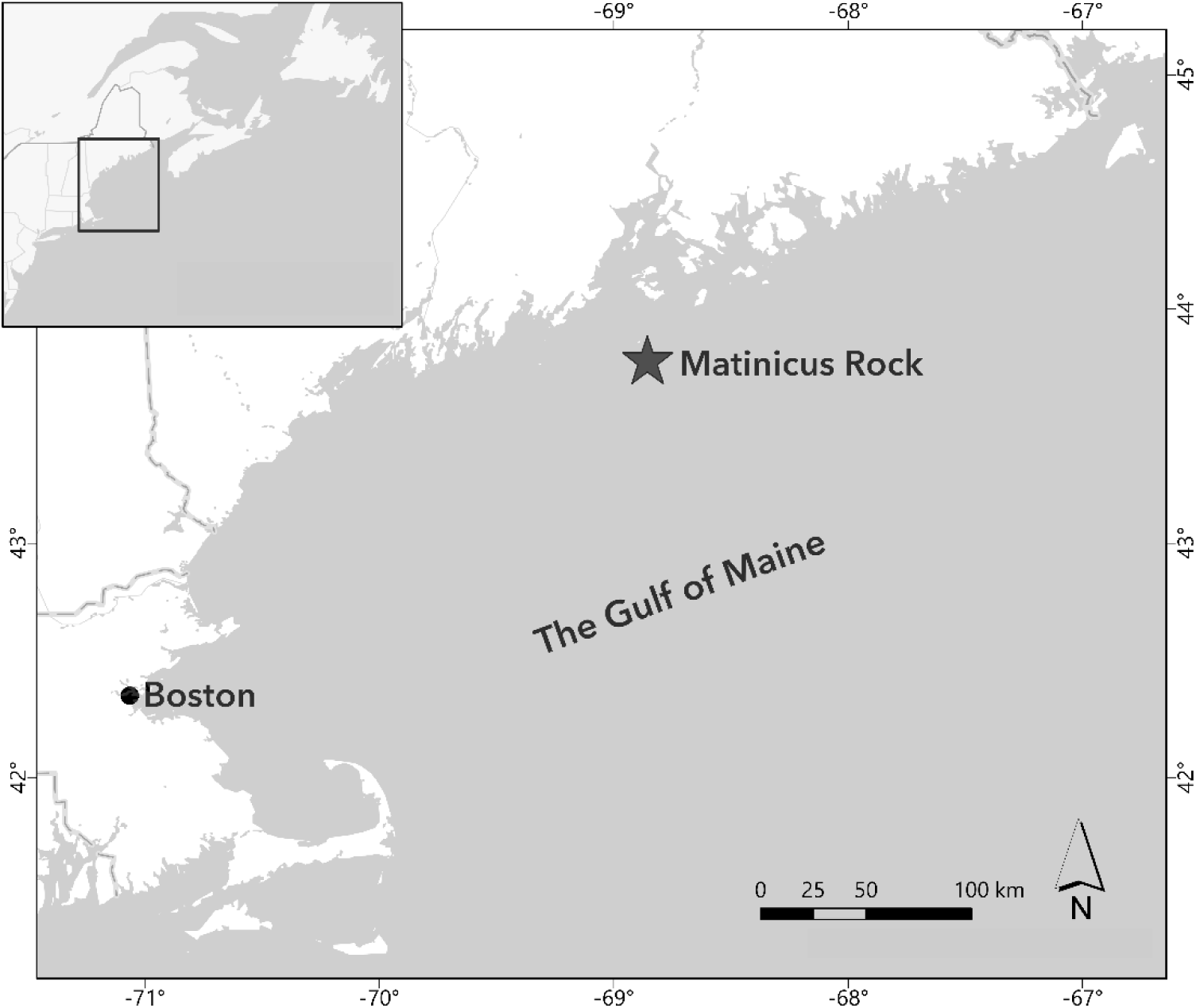
The location of the Matinicus Rock puffin colony within the Gulf of Maine.

#### 2.1.2 Field Methods

Researchers continued long-term, photography-based diet monitoring during 2021 and 2022, beginning soon after the first puffin chicks hatched (late June) and continuing until most chicks had fledged (mid August). Bill-loads of provisioning adults were photographed and the prey items in each load were identified, counted, and their sizes were estimated relative to the size of the puffin’s bill length. For more details on these photography-based diet assessments, see Kress et al. (2016) and Scopel et al. (2019).

Puffin fecal samples were collected beginning with the researchers’ arrival to the island during puffin incubation (late May/early June) and continued until most chicks had fledged (mid August). During the chick-rearing period, the timing of photography-based puffin diet observations and fecal sample collection overlapped. Fecal samples were all collected with fresh, individually-wrapped wooden spatulas, with effort made to limit contact between the spatula and adjacent surfaces. Samples were immediately placed into sterile collection vials containing 1 ml of DNA/RNA Shield (Zymo Research, Irvine, CA). Vials were stored out of direct sunlight at room temperature until they were sent for processing immediately following puffin fledging at the end of each field season (range: 1-12 weeks following sample collection).

During the incubation period, burrows known to contain an adult and egg were observed for incubation switches, indicated by one adult returning from sea and entering the burrow with the other adult typically exiting soon after. The identification of breeding individuals was facilitated by the large number of uniquely marked birds at the Matinicus Rock colony. If either of the adults defecated immediately prior to or following the incubation switch, these samples were then promptly collected. Additional incubation period samples were obtained during productivity checks when fresh feces were observed deep in burrows adjacent to adults incubating eggs. Since disturbance may cause incubating puffins to abandon their nests (Rodway et al., 1996), we did not remove any adults from their burrows for fecal sample collection.

Following chick hatch, we no longer collected samples from within burrows due to uncertainty about the origin of the fecal matter. Instead, adult fecal samples were obtained immediately before or after observed chick-provisioning events, as nonbreeding puffins attending the colony do not carry fish. We avoided collecting samples if there was doubt as to which fecal deposit in the area was made by the observed bird of known breeding status. Chick fecal samples were collected opportunistically while handling chicks for regular growth and productivity monitoring efforts. When samples were collected during the handling of chicks as part of monitoring efforts, birds were positioned over clean sheets of aluminum foil or wax paper to minimize possible environmental contamination. Fecal samples from non-restrained birds were only collected from surfaces free of visible contamination (i.e., other seabird fecal matter). To determine if amplifiable DNA existed within the local environment, field blanks were collected from varied surfaces and regions of the colony in order to confirm that our samples could be considered reasonably free of environmental contamination.

Previous research indicates that detectable quantities of prey DNA may be detected in seabird feces up to four days after the ingestion of that prey (Deagle et al., 2010). Accordingly, samples were not collected from the same individuals or burrows less than five days apart to avoid repeated sampling. Aside from avoiding particular burrows for this reason, we attempted to collect samples from puffins across the breeding colony to minimize the influence of potential sub-colony variation in diet (Hipfner et al., 2007).

#### 2.1.3 Laboratory and Bioinformatics Methods

DNA extraction, amplification, and sequencing closely followed methods outlined in Fayet et al. (2021, supplemental information) and were led by GVC. Fecal samples were homogenized and DNA was extracted from samples using a Quick-DNA Fecal/Soil Microbe Miniprep Kit (Zymo Research, Irvine, CA). We followed a hierarchical barcoding approach, targeting two gene regions using primers with different taxonomic breadth and resolution. We first used universal metazoan primers to target a region of the 18S gene to capture the occurrence of broad metazoan taxa within puffin diet (McInnes et al., 2017). Since we expected fish to be the primary component of puffin diet, we also employed MiFish primers (Miya et al., 2015) that target the 12S gene to obtain higher taxonomic resolution of the fish DNA detected in fecal samples. For both primer sets, DNA sequences were amplified using 4.6 μl of template DNA and a two-step PCR. Field-, extraction-, and PCR-blanks were used to monitor for contamination at every step. Samples and blanks were sent to the University of New Hampshire for sequencing on an Illumina HiSeq 2500 (Illumina Inc., San Diego, CA).

Following sequencing, demultiplexed reads were imported into Qiime2 (Bolyen et al., 2019) for processing (all code can be found at https://github.com/GemmaClucas/Matinicus-Rock-2021-Atlantic-Puffins/). Sequence reads for the 12S region were identified using an iterative blast method with a custom database downloaded from GenBank using the RESCRIPt plugin (Robeson et al., 2021). We removed all avian and human DNA from our data set, as well as taxonomic groups known to solely contain obligate parasites, since these did not represent intended prey. Alpha diversity rarefaction curves were constructed to determine the read depth necessary to capture the complete diversity of the community present in each sample; from these rarefaction curves, we determined that 2500 sequences per sample were required for 18S analyses and 600 sequences per sample were required for 12S analyses. Samples were rarefied to these depths and all samples with an insufficient number of sequences were removed from our data set.

Taxonomic assignments of sequences were verified using the National Center for Biotechnology Information’s BLAST tool (NCBI, 2023) and the geographic distributions of identified fish taxa were checked using FishBase (Froese and Pauly, 2023) to ensure assignments were reasonable. For 12S taxa, taxonomic assignments matching reference sequences with 98% similarity or greater were assigned to the species-level while taxa with lower percentages were assigned to the genus or family level, depending on the number of similarly-ranked, sympatric species. Due to lower possible taxonomic resolution, 18S taxonomic assignments were grouped by class or order.

### 2.2 Data Processing

#### 2.2.1 Observational Methods

Once prey items were identified and counted from photographs, the frequency of occurrence (FOO) of prey types were calculated by year. FOO data summarizes the proportion of samples in a group that contain the taxa of interest. We assumed each provisioning bill load represented a discrete sample.

Using photographs, the size of fish prey identified in bill loads could be estimated in increments of 0.25 relative to a mean puffin bill length of 30 mm (Harris and Wanless, 2012). The approximated lengths of prey items could then be used to estimate the mass of prey delivered to puffin chicks using published length-mass relationships (Winters, 1989; Wigley et al., 2003; see Supplemental Information, Table 1). Totaling our estimates of prey biomass delivered, we could then estimate the relative biomass contributed by each prey taxa per year. The importance of each prey taxa in our observation-based diet assessments could thus be summarized using FOO or estimated relative biomass. Species occurring infrequently (FOO < 5%) were excluded from these summaries.

**Table 1.** Sample sizes for diet analyses split by methodology, bird age, breeding stage, and study year. For molecular methods, these numbers represent those samples that amplified sufficient DNA to be included; of 414 total fecal samples collected, 43% of samples (n = 176) had sufficient 18S reads while 69% of samples (n = 285) had sufficient 12S reads. Note that not all methods can produce diet estimates for each age or breeding stage.

| Method | Stage | 2021 |  | 2022 |  |
| --- | --- | --- | --- | --- | --- |
|  |  | Adults | Chicks | Adults | Chicks |
| Bill-load Photography |  |  |  |  |  |
|  | Incubation | - | - | - | - |
|  | Chick-rearing | - | 324 | - | 368 |
| 18S (Broad Metazoan Taxa) |  |  |  |  |  |
|  | Incubation | 3 | - | 4 | - |
|  | Chick-rearing | 43 | 31 | 43 | 52 |
| 12S (Fish Taxa) |  |  |  |  |  |
|  | Incubation | 11 | - | 12 | - |
|  | Chick-rearing | 83 | 43 | 65 | 71 |

#### 2.2.2 Molecular Methods

Metabarcoding-based diet analyses frequently summarize the importance of different prey types using either FOO or relative read abundance (RRA; Deagle et al., 2019). RRA measures the proportion of reads within a sample or group assigned to that taxon. We relied on FOO, the more conservative measurement, for 18S data in our assessment of broad metazoan taxa in puffin diet. For a taxon to qualify as present in a fecal sample, we set a minimum threshold for each prey type of at least 1% of the samples’ total read depth (i.e., requiring at least 25 reads for a sample depth of 2500 reads) to limit the influence of rare taxa. In most cases, this removed taxa that were also detected in field blanks yet were unlikely to have been consumed by puffins, such as Diptera flies. Using this presence/absence data, we calculated the FOO of all broad metazoan taxa in different combinations of bird age, breeding stage, and study year.

For 12S (fish) data, we calculated both FOO and RRA. Previous research suggests RRA can be employed as an imperfect but useful proxy for the relative biomass of a given prey type consumed (Clucas et al., in press); RRA can thus aid in determining the relative importance of different prey types in an organism’s diet (Bowles et al., 2011; Deagle et al., 2019). This was of particular interest for fish, as fish were expected to constitute the majority of puffin diet by mass (Harris and Wanless, 2012). Using the same groupings of bird age, breeding stage, and study year as with 18S data, we calculated the relative proportion of reads contributed by each species to the grouping’s total.

Following the identification and taxonomic assignment of all sample sequences, we summarized the diversity of prey types puffins consumed by calculating the species richness for each sample in our analyses. Shannon Index (Shannon, 1948) values were then calculated for RRA and relative biomass estimates using the ‘diversity’ function in the R (Version 4.0.2; R Core Team, 2021) package ‘vegan’ (Oksanen et al., 2022). To determine if we collected a sufficient number of fecal samples to effectively characterize the diets of incubating adults, chick-rearing adults, and chicks, we constructed prey species accumulation curves using the ‘specaccum’ function in R package ‘vegan’ using 1,000 random permutations (Supplementary Material, Figure 1).

### 2.3 Statistical Analyses

#### 2.3.1 Comparing Methodology

In order to first assess the comparability of our two methods of diet assessment, we tested for differences in our estimates of puffin chick diet as obtained from bill-load photography and fecal DNA metabarcoding. For these comparisons, we utilized an abridged data set of eight dominant fish prey types that together comprised more than 95% of prey sequences and were identifiable via both methods of diet assessment. We calculated the estimated consumption of each of the main fish prey items for three, two-week windows, representing the early, middle, and late parts of the chick-rearing period. Our different estimates (FOO, RRA, relative biomass) of the relative consumption of each prey type were compared with Pearson correlation tests using the ‘cor.test’ function in the R package ‘stats’ (R Core Team, 2021).

We then tested if the overall estimated composition of puffin diets obtained from observational and molecular methods were statistically distinguishable using two tests of community similarity. Data summarized as presence-absence for FOO calculations were tested using analyses of similarity (ANOSIM) with method as a fixed effect, using the ‘anosim’ function in R package ‘vegan’ (Oksanen et al., 2022). ANOSIM is a nonparametric means of testing for significant differences between two groups using ranked dissimilarities, testing if greater differences exist between groups or within them. Following Bowser et al. (2013), a Bray-Curtis dissimilarity measure and 999 permutations were used for all ANOSIMs throughout our study.

Estimates of puffin diet composition using proportion data (relative biomass and RRA) were compared using permutational multivariate analyses of variance (PERMANOVA) using the ‘adonis2’ function, also in R package ‘vegan’ (Oksanen et al., 2022). PERMANOVA is a nonparametric alternative to MANOVA and is more robust than ANOSIM for data with heterogenous dispersions (Anderson and Walsh, 2013). We used a significance level of p < 0.05 for all PERMANOVA and ANOSIM. Following each significant test, a *post hoc* analysis of similarity percentages (SIMPER) was used to determine the contribution of each taxa to the observed dissimilarity of the groups examined.

#### 2.3.2 Comparing Ages, Breeding Stages, and Years

We likewise used ANOSIM and PERMANOVA to compare puffin diet composition among different age, breeding stage, and year combinations. For these tests, we used the complete community of fish prey taxa detected via 12S molecular methods. Due to a large number of metazoan groups identified using 18S molecular methods, we removed all but those taxa identified using both molecular and observational methods, attributing the presence of the others to secondary consumption (Bowser et al., 2013). We tested for differences in puffin diet across the two age classes of puffins examined (breeding adults and chicks), the different stages of the breeding period (incubation and chick-rearing), and the two years of the study (2021 and 2022). Additionally, for each of these combinations, we used two-sample T-tests to compare the species richness and Shannon Index values for 12S fish prey.

## 3 Results

During the 2021 and 2022 field seasons, researchers performed 250 total hours of observation-based puffin diet surveys. A total of 692 photographed provisioning loads (Table 1) were included in these analyses, representing more than 2,500 identifiable prey items. For molecular dietary assessments, researchers collected a total of 414 fecal samples from Matinicus Rock puffins. Of these, 176 (43%) amplified sufficient dietary DNA using 18S primers to be included in our assessment of broad metazoan taxa represented in puffin diet. Using 12S MiFish primers, 285 (69%) amplified to a sufficient depth to be included in our analyses of fish taxa consumed by puffins.

### 3.1 Taxa Detected in Puffin Diet

Observational methods identified six broad taxonomic groups being provisioned to chicks (Table 2). Fish occurred in more than 97% of bill loads but squid, krill/shrimp, polychaetes, amphipods, and a single comb jelly were also identified. At least 14 fish taxa were identified in puffin bill-loads across the two years (Table 3), with haddock, Atlantic saury (*Scomberesox saurus*), sandlance (*Ammodytes spp.*), and rough scad among the most commonly observed species. Unidentified hake species (Merlucciidae/Phycidae/Lotidae) were observed frequently in bill loads but could not be visually identified to species level due to the similar morphology of juvenile hakes, and so were combined as a single “hake spp.” category.

**Table 2.** The frequency of occurrence (FOO, expressed as %) of prey taxa detected in puffin fecal samples using 18S primers for 2021 and 2022. FOO for the traditional, photographic identification of prey using adult provisioning loads is displayed for comparison. Bolded taxa are those identified using both methods and most likely to represent primary consumption.

| <u>**Taxa (Common Name)**</u> | **2021** |  |  |  | **2022** |  |  |  |
| --- | --- | --- | --- | --- | --- | --- | --- | --- |
|  | <u>**INC.**</u><br>**Adults**<br>*n=3* | **Adults**<br>*n=43* | <u>**CHICK-REARING**</u><br>**Chicks**<br>*n=31* | **Photos**<br>*n = 324* | <u>**INC.**</u><br>**Adults**<br>*n = 4* | **Adults**<br>*n = 43* | **Chicks**<br>*n = 52* | **Photos**<br>*n = 368* |
| Acoela (Acoelomorph Flatworms) |  |  |  |  |  | 2.3 |  |  |
| **Actinopterygii (Ray-finned Fishes)** | **33.3** | **97.7** | **100.0** | **96.9** | **75.0** | **90.7** | **100.0** | **98.4** |
| Amiphoda (Amphipods) |  |  |  |  |  |  |  | 0.8 |
| *Indet.* Animalia (Metazoans) | 33.3 | 20.1 | 3.2 |  | 25.0 | 16.3 | 3.9 |  |
| Copepoda (Copepods) | 66.7 | 58.1 | 51.6 |  | 50.0 | 60.5 | 50.0 |  |
| Chromadorea (Nematodes) |  | 2.3 | 9.7 |  |  |  | 5.8 |  |
| Cladocera (Water Fleas) |  |  |  |  |  | 2.3 | 1.9 |  |
| Collembola (Springtails) |  |  |  |  |  |  | 1.9 |  |
| **Cydippida (Comb Jellies)** |  | **4.6** |  | **0.3** |  |  | **1.9** |  |
| Demospongiae (Most Sponges) |  |  | 6.4 |  |  | 2.4 | 3.9 |  |
| **Eucarida (Krill, Shrimp, etc.)** |  | **23.2** |  |  | **25.0** | **23.3** |  | **1.9** |
| Hydrozoa (Hydrozoans) |  | 40.0 | 35.5 |  |  | 16.3 | 13.5 |  |
| Mytiloidea (True Mussels) |  | 2.3 |  |  |  |  |  |  |
| Neoptera (Most Winged Insects) |  | 4.7 | 3.2 |  |  | 4.7 | 3.9 |  |
| **Polychaeta (Polychaetes)** | **33.3** |  | **3.2** | **0.6** |  | **4.7** |  |  |
| Scyphozoa (True Jellyfish) |  | 18.6 | 19.4 |  |  | 9.3 | 1.9 |  |
| **Teuthida (Squid)** |  | **4.6** | **3.2** | **3.7** |  | **11.6** | **1.9** | **0.8** |

Inc. Incubation stage

**Table 3.**
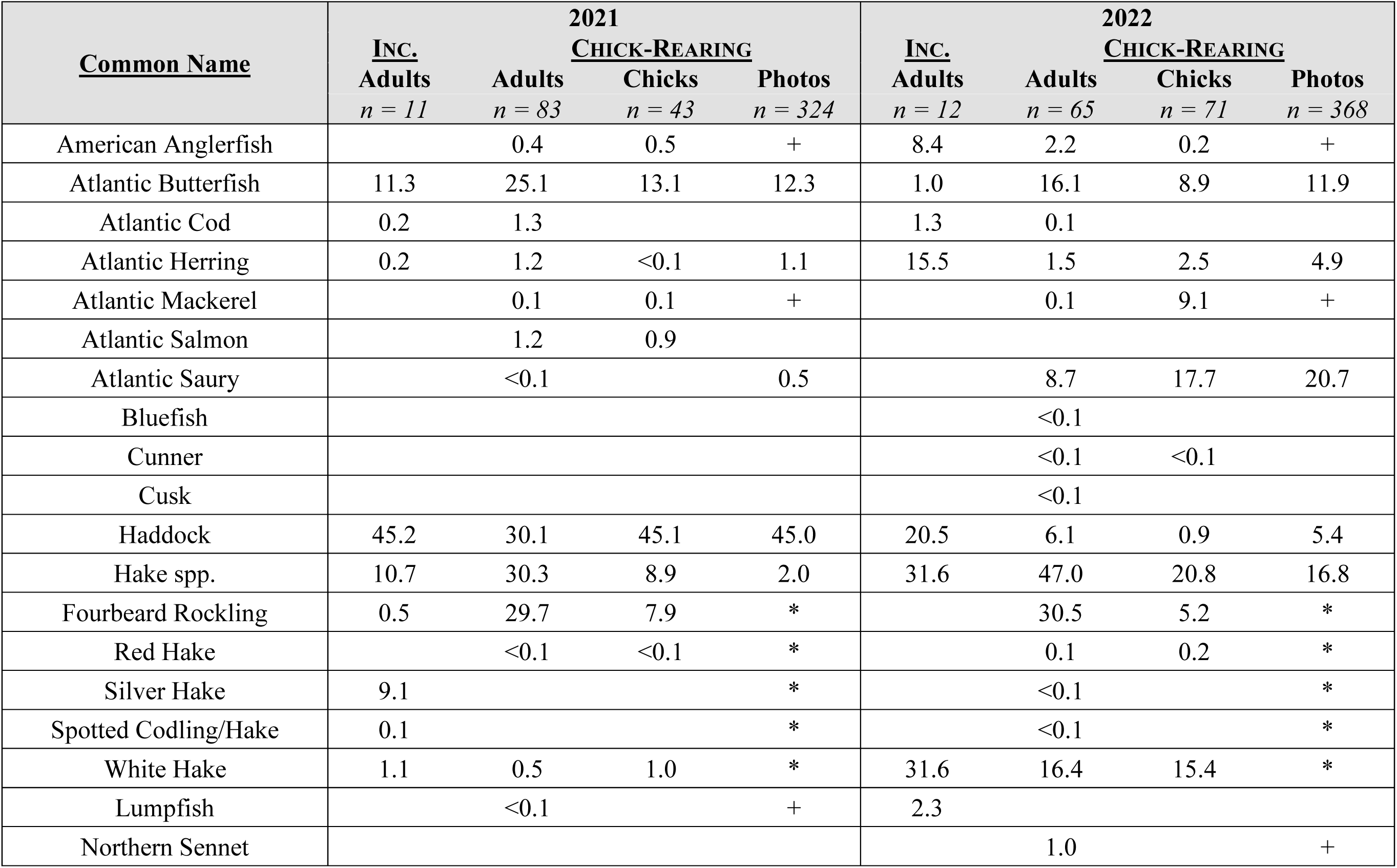
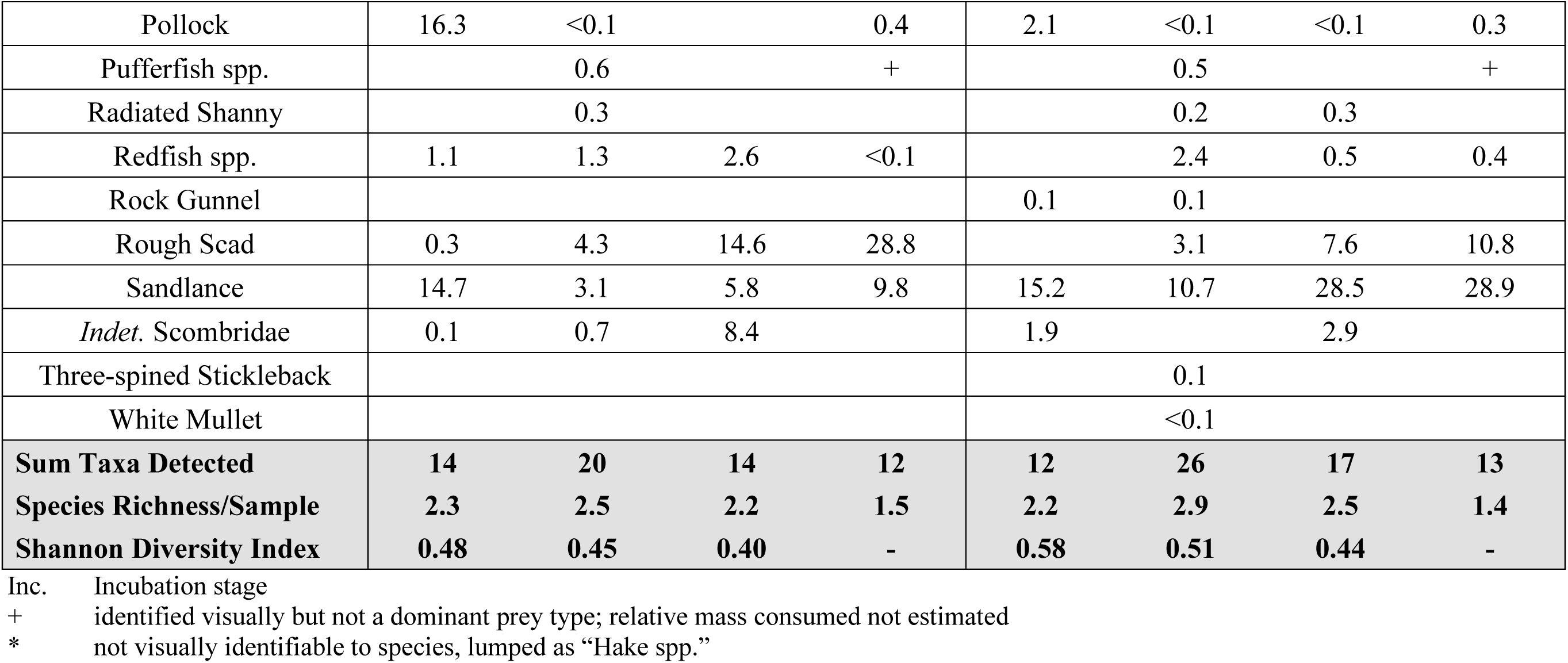
The relative read abundance (RRA, expressed as %) of fish prey taxa detected in puffin fecal samples using 12S primers for 2021 and 2022. Estimated relative biomass for the traditional, photographic identification of prey using chick provisioning loads is displayed for comparison. For scientific names of prey taxa, see Supplemental Materials, Table 2.

| <u>Common Name</u> | **2021** |  |  |  | **2022** |  |  |  |
| --- | --- | --- | --- | --- | --- | --- | --- | --- |
|  | <u>INC.</u><br>Adults<br>*n* = 11 | <u>CHICK-REARING</u><br>Adults<br>*n* = 83 | <u>CHICK-REARING</u><br>Chicks<br>*n* = 43 | <u>CHICK-REARING</u><br>Photos<br>*n* = 324 | <u>INC.</u><br>Adults<br>*n* = 12 | <u>CHICK-REARING</u><br>Adults<br>*n* = 65 | <u>CHICK-REARING</u><br>Chicks<br>*n* = 71 | <u>CHICK-REARING</u><br>Photos<br>*n* = 368 |
| American Anglerfish |  | 0.4 | 0.5 | + | 8.4 | 2.2 | 0.2 | + |
| Atlantic Butterfish | 11.3 | 25.1 | 13.1 | 12.3 | 1.0 | 16.1 | 8.9 | 11.9 |
| Atlantic Cod | 0.2 | 1.3 |  |  | 1.3 | 0.1 |  |  |
| Atlantic Herring | 0.2 | 1.2 | <0.1 | 1.1 | 15.5 | 1.5 | 2.5 | 4.9 |
| Atlantic Mackerel |  | 0.1 | 0.1 | + |  | 0.1 | 9.1 | + |
| Atlantic Salmon |  | 1.2 | 0.9 |  |  |  |  |  |
| Atlantic Saury |  | <0.1 |  | 0.5 |  | 8.7 | 17.7 | 20.7 |
| Bluefish |  |  |  |  |  | <0.1 |  |  |
| Cunner |  |  |  |  |  | <0.1 | <0.1 |  |
| Cusk |  |  |  |  |  | <0.1 |  |  |
| Haddock | 45.2 | 30.1 | 45.1 | 45.0 | 20.5 | 6.1 | 0.9 | 5.4 |
| Hake spp. | 10.7 | 30.3 | 8.9 | 2.0 | 31.6 | 47.0 | 20.8 | 16.8 |
| Fourbeard Rockling | 0.5 | 29.7 | 7.9 | * |  | 30.5 | 5.2 | * |
| Red Hake |  | <0.1 | <0.1 | * |  | 0.1 | 0.2 | * |
| Silver Hake | 9.1 |  |  | * |  | <0.1 |  | * |
| Spotted Codling/Hake | 0.1 |  |  | * |  | <0.1 |  | * |
| White Hake | 1.1 | 0.5 | 1.0 | * | 31.6 | 16.4 | 15.4 | * |
| Lumpfish |  | <0.1 |  | + | 2.3 |  |  |  |
| Northern Sennet |  |  |  |  |  | 1.0 |  | + |
| Pollock | 16.3 | <0.1 |  | 0.4 | 2.1 | <0.1 | <0.1 | 0.3 |
| Pufferfish spp. |  | 0.6 |  | + |  | 0.5 |  | + |
| Radiated Shanny |  | 0.3 |  |  |  | 0.2 | 0.3 |  |
| Redfish spp. | 1.1 | 1.3 | 2.6 | <0.1 |  | 2.4 | 0.5 | 0.4 |
| Rock Gunnel |  |  |  |  | 0.1 | 0.1 |  |  |
| Rough Scad | 0.3 | 4.3 | 14.6 | 28.8 |  | 3.1 | 7.6 | 10.8 |
| Sandlance | 14.7 | 3.1 | 5.8 | 9.8 | 15.2 | 10.7 | 28.5 | 28.9 |
| *Indet.* Scombridae | 0.1 | 0.7 | 8.4 |  | 1.9 |  | 2.9 |  |
| Three-spined Stickleback |  |  |  |  |  | 0.1 |  |  |
| White Mullet |  |  |  |  |  | <0.1 |  |  |
| **Sum Taxa Detected** | **14** | **20** | **14** | **12** | **12** | **26** | **17** | **13** |
| **Species Richness/Sample** | **2.3** | **2.5** | **2.2** | **1.5** | **2.2** | **2.9** | **2.5** | **1.4** |
| **Shannon Diversity Index** | **0.48** | **0.45** | **0.40** | **-** | **0.58** | **0.51** | **0.44** | **-** |

Inc. Incubation stage + identified visually but not a dominant prey type; relative mass consumed not estimated \* not visually identifiable to species, lumped as "Hake spp."

Molecular methods identified 17 broad groups of metazoans from DNA in puffin fecal samples (Table 2). Many of these taxa likely represent secondary consumption but we assumed that the five major groups detected via both methods (fish, squid, krill/shrimp, polychaetes, and comb jellies) represent the range of metazoans groups most likely to have been consumed directly by puffins (Figure 3). The other groups detected (e.g., copepods) were likely consumed by puffin prey taxa (Bowser et al., 2013). Using 12S primers, we identified 28 unique fish taxa occurring in puffin diet (Table 3), twice the number identified via observational methods. In part, this was driven by the higher taxonomic resolution offered by DNA metabarcoding; we determined that at least five species of “hake spp.” were consumed, with white hake (*Urophycis tenuis)* and fourbeard rockling (*Enchelyopus cimbrius*) the most frequently detected. We also detected four species not previously documented in puffin diet (spotted hake *Urophycis regia*, Atlantic salmon *Salmo salar*, northern sennet *Sphyraena borealis*, and white mullet *Mugil curema*).

### 3.2 Metabarcoding as a Method for Estimating Seabird Diet

We observed a positive and significant correlation between our estimates of relative prey consumption as determined by observational and molecular means (Figure 2). Both FOO and RRA produced estimates of puffin chick diet that were strongly correlated with our estimates of relative biomass consumed from observational methods. The relationship of estimated relative biomass and RRA was stronger than that with FOO (RRA: Pearson’s r = 0.847, p < 0.001; FOO: Pearson’s r = 0.801, p < 0.001), suggesting that RRA, in particular, can serve as a useful proxy for the relative biomass of each fish prey species consumed. Accordingly, we did not detect a significant effect of methodology on the estimated composition of puffin chick diet (PERMANOVA, F_1,769_ = 1.722, p = 0.12).

**Figure 2.**
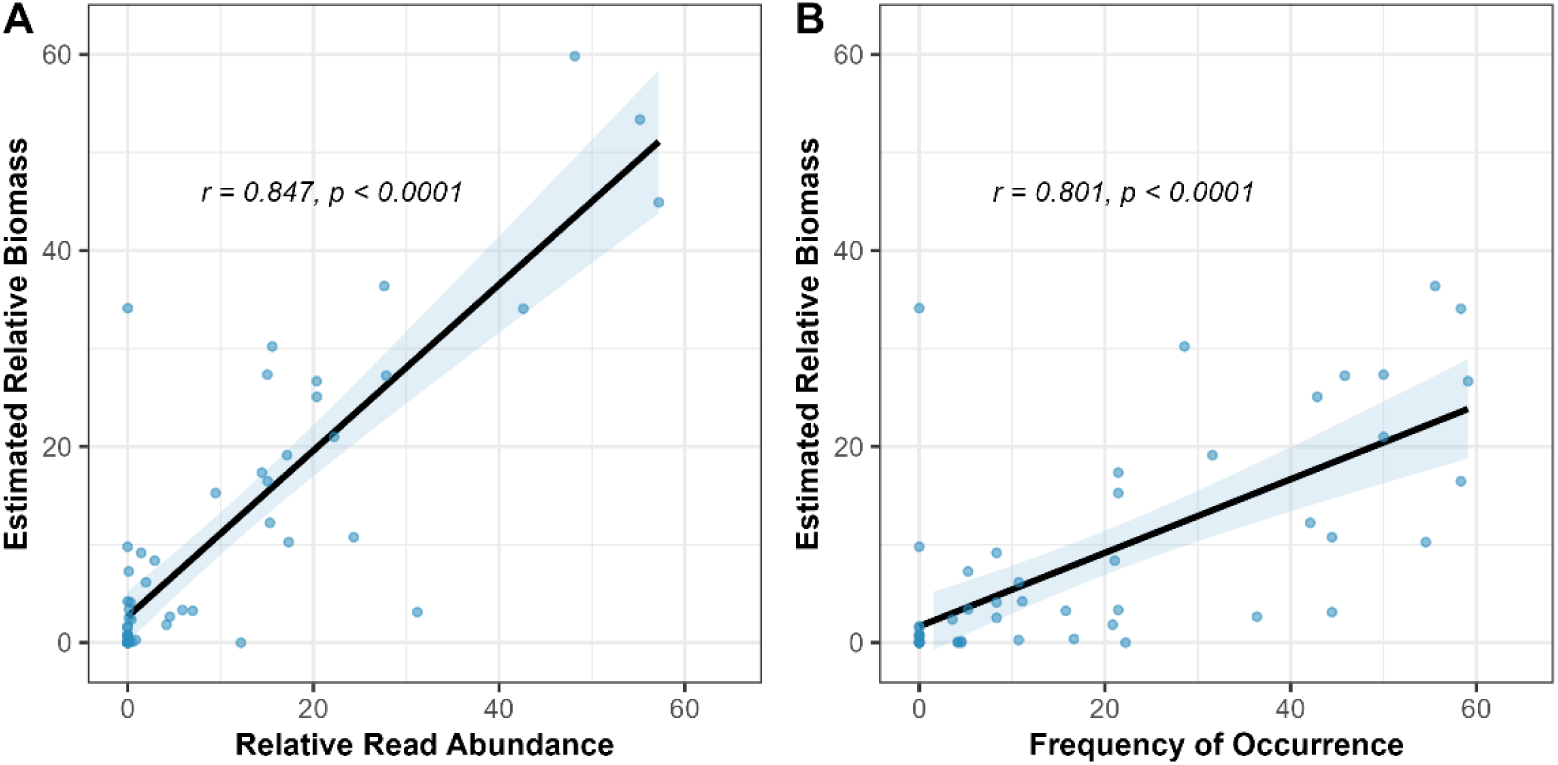
The relationships between two quantifications of DNA-assessed diet composition (RRA and FOO) and the visually-estimated relative biomass of prey taxa delivered to chicks during the same time period. Each point represents the estimated contribution of a taxa to puffin chick diet during one period (early, middle, or late) of the chick-rearing stage.

### 3.3 Variation in Puffin Diet

#### 3.3.1 Between Ages

Of the 28 unique fish taxa that occurred in our study of puffin diet overall, all 28 species were detected in adult puffin fecal samples while 17 were detected in chick samples. Mean species richness per sample was significantly higher in adult fecal samples than those from chicks during the chick-rearing period (mean of 3.1 and 2.7 species per sample, respectively; Student’s T-test, t = 2.34, df = 257.59, p = 0.02). However, Shannon Index values calculated from RRA values – therefore accounting for the evenness of taxa within a sample – did not suggest higher dietary diversity for adults than chicks (Student’s T-test, t = 1.13, df = 252.61, p = 0.26).

We detected significant differences in the occurrence of both broad prey groups (ANOSIM, R = 0.052, p < 0.001) and fish prey taxa (ANOSIM, R = 0.085, p < 0.001) between adult and chick diets using metabarcoding methods. *Post hoc* SIMPER analyses revealed that differences were driven primarily by the higher occurrence of invertebrate prey (particularly Eucarida and Teuthida) in adult diet and a higher occurrence of fish prey in chick diet (Figure 3). Similarly, we more frequently detected Atlantic saury and rough scad in chick samples while butterfish (*Peprilus triacanthus*) and hakes occurred more often in adult diet (Figure 4). The relative abundance of different fish taxa in adult and chick diets also differed, with a one-way PERMANOVA revealing significant effects of age (F_1,260_ = 11.797, p < 0.001) on diet composition. A SIMPER analysis revealed that these differences were driven largely by sandlance, rough scad, and Atlantic saury, all of which were more prevalent in chick diet.

**Figure 3.**
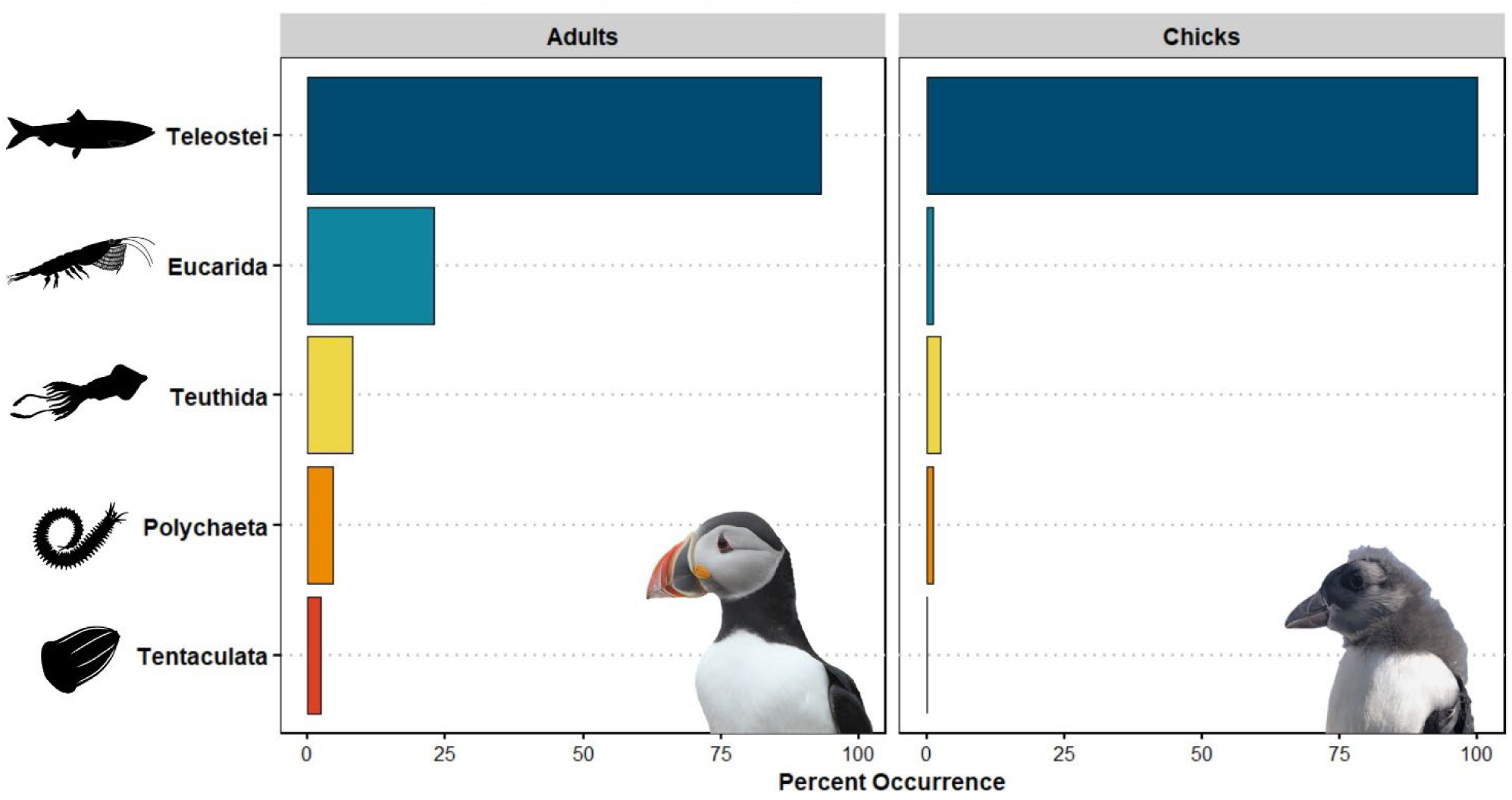
Comparison of chick diet over the two years of the study assessed via molecular methods (“DNA”, using RRA) or photography-based ones (“Obs”, estimated relative biomass).

**Figure 4.**
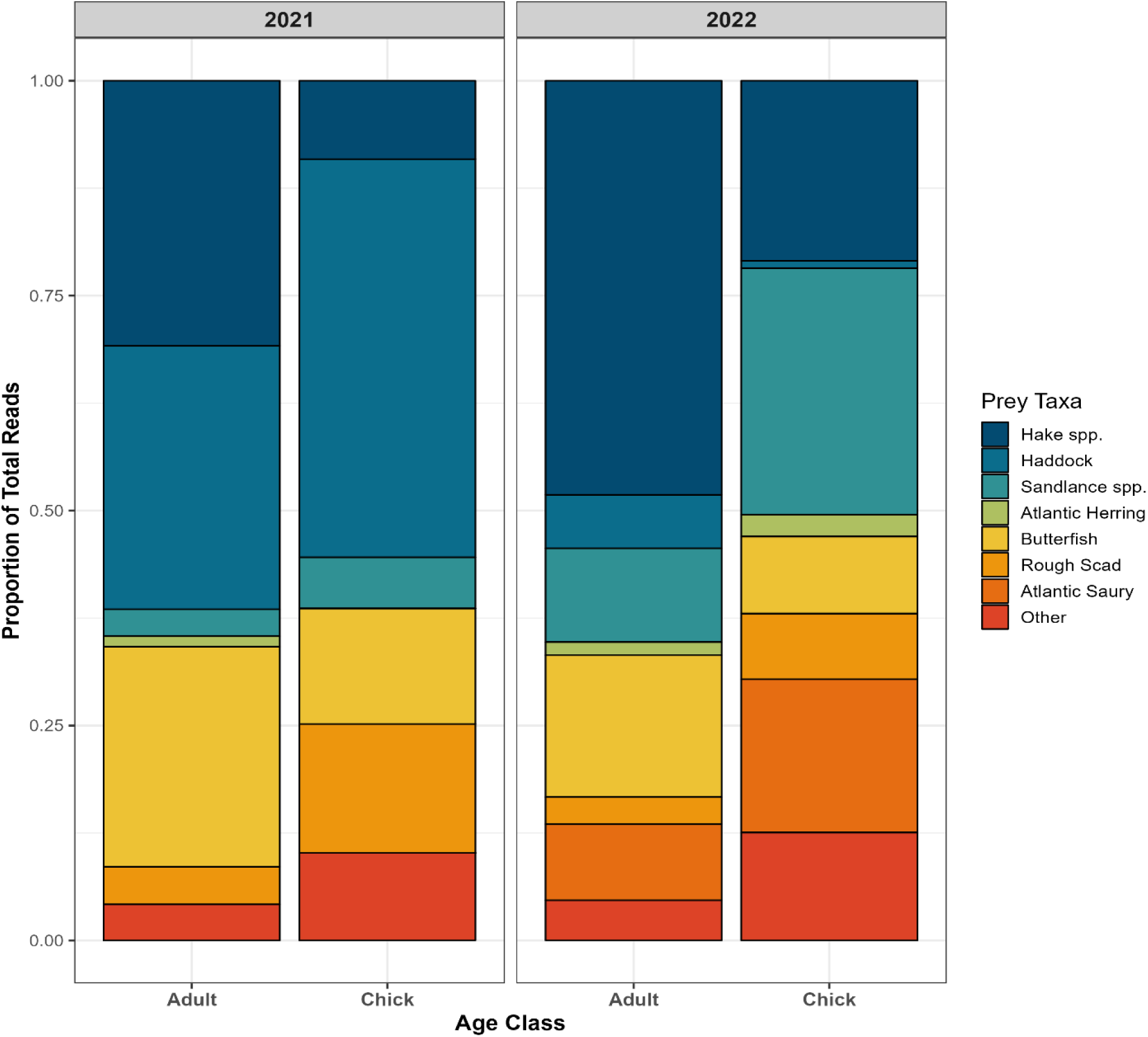
The relative read abundance of dominant fish taxa identified in puffin diet using 12S gene sequences. Data shown are only from the chick-rearing period.

#### 3.3.2 Between Stages

Amplification success was low for incubation stage fecal samples, with only 9% of these samples amplifying sufficient DNA using 18S primers to be included in analyses. Due to the very small sample sizes from the incubation period (2021: *n* = 3, 2022: *n* = 4), we could not statistically test for differences in the occurrence of broad metazoan groups. Visual examination of the data (Table 2), however, reveals evidence suggestive of less frequent fish consumption by adults during the incubation period (FOO: 57%) than during chick-rearing (FOO: 94%).

Higher amplification success for incubation stage samples using 12S primers (29% amplification success, *n* = 23 samples) allowed for comparisons of fish consumption between breeding stages (Table 3). Neither fish species richness per sample (Student’s T-test, t = 0.27, df = 26.01, p = 0.79) nor Shannon Index values (Student’s T-test, t = -0.23, df = 26.80, p = 0.82) varied with breeding stage. In contrast, the occurrence (ANOSIM, R = 0.249, p < 0.001) and relative composition (PERMANOVA, F_1,171_ = 7.01, p < 0.001) of fish taxa in adult diets did differ between between stages. A *post hoc* SIMPER analysis suggests the observed dissimilarity among fish prey was largely due to the greater consumption of Atlantic herring, white hake, and pollock (*Pollachius virens*) during the incubation period.

#### 3.3.3 Between Years

Since we determined that aspects of puffin diet varied with age class, we analyzed puffin adult and chick diets separately. The occurrence of broad metazoan taxa in puffin diet did not vary by year in either adults (ANOSIM, R = -0.08, p = 0.81) or chicks (ANOSIM, R = 0.0106, p = 0.47). Visual examination of the data confirms that the high occurrence of fish in puffin diet (especially chicks) was consistent across study years (Table 2). Similarly, greater invertebrate consumption by adults than chicks occurred in both 2021 and 2022.

In contrast, there was high interannual variability in the types of fish consumed. Within adult diet, there was significant variation between years in both the occurrence (ANOSIM, R = 0.180, p < 0.001) and relative consumption (PERMANOVA, F_1,147_ = 8.105, p < 0.001) of different fish taxa. Chick diet was also different between years (ANOSIM, R = 0.233, p < 0.001; PERMANOVA, F_1,112_ = 15.334, p < 0.001). For both age groups, the greater consumption of white hake and Atlantic saury in 2022, paired with the absence of normally-abundant haddock that year, contributed most to the observed interannual differences in prey consumption (Figure 4). Despite these differences, we did not identify any significant variation in diet diversity (species richness and Shannon Index) between years for either chicks or adults (Student’s T-tests, p ≥ 0.05).

## 4 Discussion

Using a combination of observational and molecular methods, we demonstrate significant variation in Atlantic puffin diet across ages, breeding stages, and years. This study provides the first formal assessment of puffin diet during the incubation period and is the first multi-year comparison of breeding puffin adult and chick diets using fecal DNA metabarcoding. Emerging, molecular methods produced similar estimates of chick prey consumption as traditional, observational methods for dominant prey types, yet metabarcoding identified twice as many fish prey species overall. Our study reveals that puffin diet in the GoM is both diverse and highly variable.

The use of molecular methods enabled us to examine puffin diet during the incubation period for the first time. We hypothesized that, lacking the demands of frequent chick-provisioning, adult puffins would be able to perform longer, more distant foraging trips during incubation. This could enable them to access prey resources not exploited during chick-rearing, similar to the strategy used by some thick-billed murres (*Uria lomvia;* Ito et al., 2010). While we did observe significant differences between diet composition between the incubation and chick-rearing stages, an examination of the taxa driving the observed dissimilarity suggests that these differences may have more to do with changes in local prey availability than different habitat selection. In 2022, for example, the RRA of both white hake and Atlantic herring was higher during the incubation stage than during chick-rearing. Both of these species are comparatively cold water adapted (Rose, 2005; Kleisner et al., 2017) and perhaps followed suitable thermal habitat away from the breeding colony as local waters exceeded key temperature thresholds.

Other differences between periods may be explained by interspecific variation in the timing of fish transitioning from larval stages to the age-0 juveniles generally consumed by puffins. It must be noted that our sample sizes during incubation were small and were likely insufficient to detect all prey consumed (Supplementary Material, Figure 1) so diet during incubation may be more diverse than we report here. Further work on seabird diets immediately before and after egg laying is encouraged since the prey consumed during this time may have important impacts on reproductive success (Barrett et al., 2012).

Better known is how seabird diet during chick-rearing can influence reproductive output. Yet, comparatively few published studies have compared seabird adult and chick diets simultaneously (Wilson et al., 2004; Bowser et al., 2013; Fayet et al., 2021). Our results agree with the available literature in determining that the dominant prey types consumed by adults and chicks generally overlapped, but that adults consumed a greater variety of fish prey species. On an individual sample level, samples from adults had a greater mean species richness than those from chicks, although adults’ Shannon Index values were similar to those from chick samples. This may mean that adults “snacked” on more diverse prey types than chicks but generally relied on a similar number of species. While we found adult puffin diet to be highly diverse (28 fish taxa detected), the eight most common fish taxa comprised more than 90% of puffin diet in each year for both age classes.

An important finding of our study, however, is that adult and chick diets were statistically distinguishable; despite feeding on the same dominant prey types, the relative importance of these fishes in puffin diet varied by age class. Interestingly, the only previous comparison of puffin adult and chick diets in the GoM found their prey consumption to be similar (Bowser et al., 2013). However, the degree of dietary differentiation among age classes likely varies with prey conditions (Baird, 1991). Bowser et al.’s work occurred in a year (2009) of anomalously high herring abundance where adult and chick diets differed little because herring was detected in all samples. Likewise, in a multi-colony study of puffin foraging ecology (Fayet et al., 2021), locations where prey conditions and reproductive success were high (e.g., Wales) had highly similar adult and chick diets. During years of poor prey availability, as in our study, adults must be additionally selective about the prey items they provision to chicks. Greater selectivity at these times likely leads to greater differences in chick and adult diets (Burke and Montevecchi, 2009). This suggests that the use of chick diet as a proxy for adult diet could be acceptable in years of very high prey availability, but that diets may diverge greatly when prey conditions are less favorable.

As predicted by optimal foraging theory (Orians and Pearson, 1979), we found that adults appeared to prioritize higher-quality prey for chick-provisioning and fed on lower quality items themselves. Those species consumed more often by chicks, like sandlance, Atlantic saury, and mackerel (*Scomber scombrus*), were generally lipid-rich (Harris and Hislop, 1978), large (often two or more bill lengths), or both. In contrast, adults consumed more small and low-lipid prey types like juvenile hakes and invertebrates, including squid and shrimp/krill (Spitz et al., 2010). Haddock, although lipid-poor like most gadids (Harris and Hislop, 1978), were a larger component of chick diet in 2021, perhaps because they tended to be large and their mass likely compensated for a low energy density. While puffin chick diet can be an effective indicator of fish stocks in the GoM (Depot et al., 2020), we believe that, owing to potentially varying levels of selectivity by adults, the additional incorporation of adult diet data could improve our use of puffins to detect or predict variation in forage fish stocks.

Relative to previous years of observational chick diet monitoring in the GoM (Kress et al., 2016; Scopel et al., 2019), the composition of puffin diet during the two years of this study was unusual. Haddock, consistently present in GoM puffin diet since it was first noted in 2009 (Bowser et al., 2013) was exceptionally scarce during 2022. Additionally, two typically uncommon species – rough scad and Atlantic saury – were provisioned in frequencies never before recorded at this colony (National Audubon Society’s Seabird Institute, unpublished data). Both of these species favor warm waters (Collette and Klein-MacPhee, 2002) and may have followed rising temperatures into the GoM during the two anomalously warm years of our study (Mills et al., 2023). As the occurrence of herring in puffin diet has declined in association with warming ocean temperatures (Kress et al., 2016; Scopel et al., 2019), puffins in the GoM have demonstrated remarkable dietary flexibility; here, puffins appeared to exploit the infrequent, but opportune abundance of the normally-uncommon scad and saury. As generalist predators, puffins are capable of feeding on alternative prey types and diversifying their foraging strategies in order to mediate the affects of variable prey availability (Baillie et al., 2004; Schoen et al., 2018). The below-average reproductive success during this study (2004-2020 mean: 0.65 chicks nest^-1^; 2021-22 mean: 0.45; National Audubon Society’s Seabird Institute, unpublished data), however, suggests that even the exploitation of these alternative prey types was insufficient to fully compensate for the poor prey conditions during the 2021-2022 MHWs.

In addition to the unprecedented abundance of Atlantic saury and rough scad in puffin diet, this study documents the occurrence of four species in puffin diet for the first time: Atlantic salmon, spotted hake, white mullet, and northern sennet. The appearance of salmon may be related to especially high returns of spawning adults in 2020 following Penobscot River dam removals (NOAA Fisheries - News, 2020) and the occurrence of salmon in the diet of two 2021 puffins is well-timed with the spring seaward movements of smolts spawned the previous fall (Kocik et al., 2009). The presence of the other species is likely related to warming ocean conditions and shifting thermal habitat. These three taxa are typically uncommon north of Cape Cod, being more abundant within the warmer Mid-Atlantic Bight (Collette and Klein-MacPhee, 2002).

Warming conditions – exacerbated by MHWs – have led to range shifts for many marine species and the general tropicalization of the U.S. Northeast Shelf (Kleisner et al., 2017; Friedland et al., 2020a). Our detection of these novel species in puffin diet demonstrates the potential efficacy of using metabarcoding techniques and generalist, marine predators to qualitatively monitor changing marine communities in response to global climate change (Sydeman et al., 2017; Lenoir et al., 2020).

The greater taxonomic resolution offered by metabarcoding also enabled us to identify more fish taxa in puffin fecal samples than with visual observations alone. Precise visual identification of juvenile hakes from bill load photography is not possible, and observations must be grouped as “hake spp.” in these assessments. In contrast, we were able to reliably identify five unique species of “hakes” (fourbeard rockling and white, red (*Urophycis chuss*), silver (*Merluccius bilinearis*), and spotted hakes) using fecal DNA. With molecular identification, we observed changes in the composition of “hake spp.” over our study; whereas white hake predominated during incubation stages and throughout 2022, hakes in 2021 were overwhelmingly fourbeard rockling. While seemingly trivial, closely related species may have different thermal tolerances and responses to changing ocean conditions (Rose, 2005). Morphologically-similar species can also have different lipid contents, resulting in variable energetic gain for their predators (Spitz et al., 2010). The use of diet assessment methods capable of such high-level identification can thus be valuable for understanding the complex relationships between predator diet, reproduction, and ocean warming.

Despite these advantages, metabarcoding techniques are not a panacea and cannot replace certain types of data collected using traditional methods. Most obviously, metabarcoding is not capable of estimating the total mass of prey consumed or the size or age class distributions of prey items. Whereas stomach flushing revealed that common murre (*Uria aalge*) adults tend to consume smaller, younger age classes of sandeels (*Ammodytes spp.*) than they provision their chicks (Wilson et al., 2004), the isolated use of molecular methods would be unable to account for this. Furthermore, some interpretation of molecular diet assessments are necessary to account for potential secondary and incidental ingestion (Sheppard et al., 2005). We detected the occurrence of 17 unique metazoan groups in puffin diet but this total includes many taxa unlikely to have been consumed directly by puffins due to habitat (e.g., terrestrial winged insects) or size (e.g., copepods). We thus relied heavily on concurrent visual observations of prey deliveries to determine which invertebrate taxa were most likely consumed by puffins directly and which were not. Since we detected various differences in adult and chick diets, this method is likely imperfect and even our dual-methodology here may be inadequate to fully assess adult puffin invertebrate consumption. Although our decisions regarding which taxa represent intentional consumption were largely substantiated by the available literature (Piatt, 1987; Falk et al., 1992; Bowser et al., 2013; Harris et al., 2015), it’s possible that adult puffins exploit invertebrate prey resources more broadly than currently thought.

Generally, though, we believe our dual-method strategy for assessing puffin diet confirms the utility of fecal DNA in supplementing traditional methods of diet assessment. A key finding of our research is that the estimated relative contribution of a prey taxa to puffin diet was found to be strongly correlated when estimated through traditional and molecular methods, as confirmed recently elsewhere (Clucas et al., 2024). As in other metabarcoding studies, the correlation was strong though imperfect due to various factors such as the differential survival of DNA during digestion, interspecific variation in the amount of DNA per gram of tissue, primer binding biases, and PCR stochasticity (Deagle and Tollit, 2007; Thomas et al., 2016; Alberdi et al., 2019). Despite this, the estimated relative importance of most dominant prey types was similar between methods and we believe this highlights the capability of using metabarcoding to quantitively assess seabird diet. We believe the relative composition of puffin diet may be reliably estimated using the RRA from fecal DNA metabarcoding methods, including for ages and breeding stages not previously possible to assess noninvasively.

Further monitoring of puffin diet within the GoM will provide valuable information on how puffins are adapting to changing prey-resources. The loss of puffins’ dominant prey species in other regions has very rapidly led to widespread breeding failure (Barrett et al., 1987; Miles et al., 2015) but declines in reproductive success within the GoM have so far been milder (Scopel et al., 2019). The high diversity of prey taxa available in the region has likely served as a buffer against the decline of herring and other cold water-adapted species (the portfolio concept; Schindler et al., 2015). This diversity and the opportunistic nature of puffins in exploiting irregularly-occurring prey resources may help this population be resilient to ongoing and projected oceanographic changes within the GoM.

## Acknowledgments

We are indebted to the National Audubon Society’s Seabird Institute’s for providing funding and valuable logistical support. We are thankful for the support and encouragement of the U.S. Fish & Wildlife Service and the Maine Coastal Islands NWR staff. Invaluable assistance with fieldwork was provided by Tracey Faber, Ayla Liss, Molly Henling, Keenan Yakola, Emily Onderbeke, Alyssa Eby, and Ryan Mong. We thank John Drury for his excellent seamanship and for consistently providing safe transport to Matinicus Rock aboard the *Skua*. We also thank Bronwyn Butcher and Brian Trevelline for their help and advice regarding lab work. This work constitutes part of William L Kennerley’s MSc thesis, Oregon State University.

## Supplemental Information

**Supplemental Table 1.** Length-weight relationships used to estimate the relative biomass of different prey groups appearing in chick diet, as assessed via photography-based studies. ‘Length’ is the estimated size of a prey item, measured in puffin bill lengths to the nearest 0.25 by the observer. ‘BL’ is a constant representing the estimated length of a puffin bill in centimeters (3.0 cm; Harris & Wanless, 2012). ‘Length’ and ‘BL’ are multiplied to obtain an estimate of prey length in centimeters. Estimated mass is in grams. The formula for long-bodied fish is from Winters (1989), all others are from Wigley et al. (2003).

|  |  |
| --- | --- |
| **Atlantic Herring** | $Mass = (2.71828^{-11.7972+2.9794*\ln(Length*BL)}) * 1000$ |
| **Hake spp.**<br>*Red, White, Silver, and Spotted Hakes,*<br>*Fourbeard Rockling* | $Mass = (2.71828^{-12.95+3.2903*\ln(Length*BL)}) * 1000$ |
| **Haddock and allies**<br>*Gadidae spp.* | $Mass = (2.71828^{-11.8062+3.0766*\ln(Length*BL)}) * 1000$ |
| **Deep-bodied Fish**<br>*Butterfish, Rough Scad* | $Mass = (2.71828^{-11.6824+3.293*\ln(Length*BL)}) * 1000$ |
| **Long-bodied Fish**<br>*Sandlance, Atlantic Saury, etc.* | $Mass = 0.0000015 * (Length * BL)^{3.13}$ |

**Supplemental Table 2.** Comprehensive list of all fish species detected in Atlantic puffin diet on Matinicus Rock, 2021-2022.

| **<u>Order</u>** | **<u>Family</u>** | **<u>Scientific Name</u>** | **<u>Common Name</u>** |
| --- | --- | --- | --- |
| Beloniformes | Scomberesocidae | *Scomberesox saurus* | Atlantic Saury |
| Carangiformes | Carangidae | *Trachurus lathami* | Rough Scad |
| Clupeiformes | Clupeidae | *Clupea harengus* | Atlantic Herring |
| Gadiformes | Gadidae | *Gadus morhua* | Atlantic Cod |
|  |  | *Melanogrammus aeglefinnus* | Haddock |
|  |  | *Pollachius virens* | Pollock |
|  | Lotidae | *Brosme brosme* | Cusk |
|  |  | *Enchelyopus cimbrius* | Fourbeard Rockling |
|  | Merlucciidae | *Merluccius bilinearis* | Silver Hake |
|  | Phycidae | *Urophycis chuss* | Red Hake |
|  |  | *Urophycis regia* | Spotted Codling/Hake |
|  |  | *Urophycis tenuis* | White Hake |
| Istiophoriformes | Sphyraenidae | *Sphyraena borealis* | Northern Sennet |
| Labriformes | Labridae | *Tautogolabrus adspersus* | Cunner |
| Lophiiformes | Lophiidae | *Lophius americanus* | American Anglerfish |
| Mugiliformes | Mugilidae | *Mugil carema* | White Mullet |
| Perciformes | Pomatomidae | *Pomatomus saltatrix* | Bluefish |
| Salmoniformes | Salmonidae | *Salmo salar* | Atlantic Salmon |
| Scombriformes | Scombridae | *Scomber scombrus* | Atlantic Mackerel |
|  | Stromateidae | *Peprilus triacanthus* | Atlantic Butterfish |
| Scorpaeniformes | Cyclopteridae | *Cyclopterus lumpus* | Lumpfish |
|  | Gasterosteidae | *Gasterosteus aculeatus* | Three-spined Stickleback |
|  | Pholidae | *Pholis gunnellus* | Rock Gunnel |
|  | Scorpaenidae | *Sebastes spp.* | Redfish spp. |
|  | Stichaeidae | *Ulvaria subbifurcata* | Radiated Shanny |
| Tetraodontiformes | Tetraodontidae | *Sphoeroides spp.* | Pufferfish spp. |
| Trachiniformes | Ammodytidae | *Ammodytes spp.* | Sandlance |

**Supplemental Figure 1.**
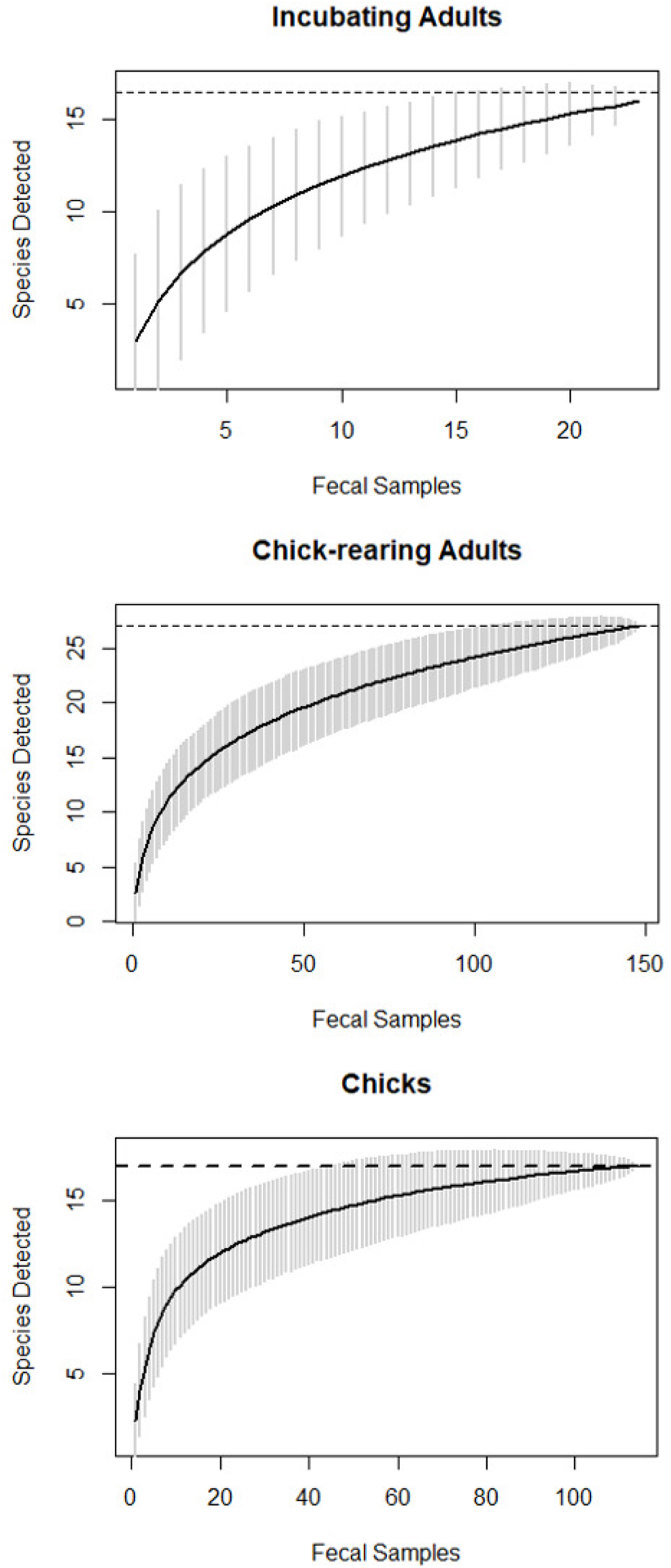
Species accumulation curves showing the estimated increase in fish prey species detected per age group and breeding stage for each additional fecal sample.

**Supplemental Figure 2.**
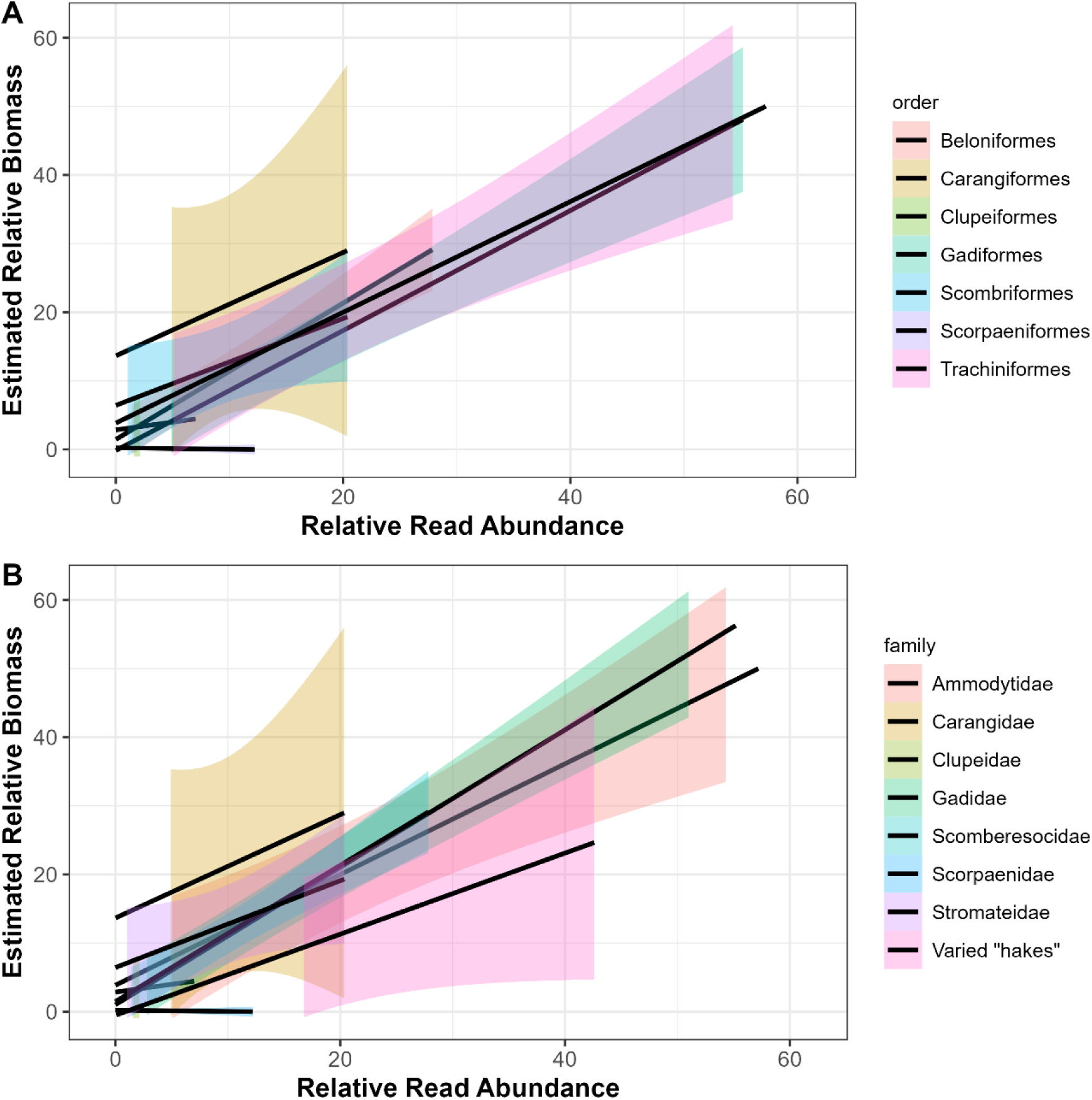
Correlations between the relative read abundance and estimated relative biomass of different prey taxa in Atlantic puffin chick diet by taxonomic order and family.

**Figure 3.**
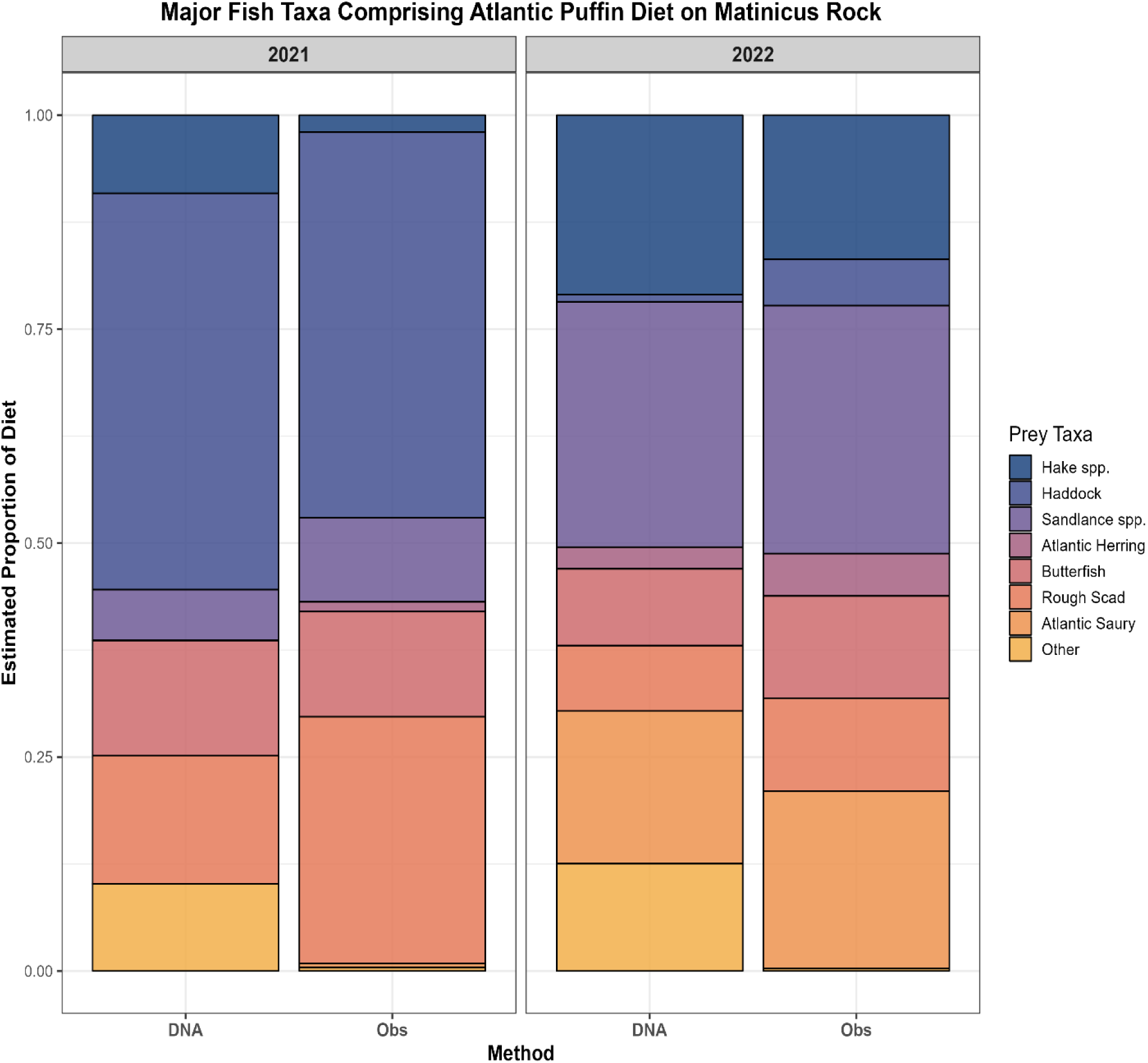
The frequency of occurrence of major metazoan taxa identified using 18S gene sequences in puffin adult and chick diet. Taxa shown are those that likely represent primary consumption. Data from the two years of the study were combined after it was found that no significant differences existed between them.

